# Hō‘ike: A Joint-Embedding Predictive Architecture for Transcriptome Data Generation with Diffusion Models

**DOI:** 10.64898/2026.08.03.741845

**Authors:** Phillip Souza, Colby T. Ford

## Abstract

In biomarker discovery, access to sufficient quantities of condition-specific transcriptomic data is often limited by cohort size, privacy concerns, and domain shift between normal and condition populations. Generative modeling can augment scarce cohorts and probe distributional transitions. Furthermore, synthetic transcriptome generation can support differential expression analyses, machine learning, privacy-preserving data sharing, benchmarking, and hypothesis generation in translational bioinformatics workloads in fields such as oncology.

Here, we present *Hoike*, a framework that combines a crossdomain Joint-Embedding Predictive Architecture (JEPA) with a latent diffusion model to generate condition-specific bulk transcriptomes from a normal reference context. In *Hoike*, normal tissue profiles provide continuous conditioning signals, while the model learns disease-linked shifts in latent space and reconstructs gene-level expression in log_2_(TPM+1) space. The implementation supports paired normal-condition training, tissue-aligned conditioning, and constrained non-negative decoding for biologically valid outputs. We describe the architecture, objective design, and evaluation protocol used in this work across GTEx-derived normal references and multiple TCGA condition cohorts as a case study. This serves as the technical specification of the *Hoike* framework and its reproducible analysis workflow.

## Introduction

Bulk RNA-sequencing studies have enabled foundational atlases of normal and condition-associated transcriptional states, including large tissue-resolved reference resources from GTEx (1) and cancer-focused cohorts from The Cancer Genome Atlas (TCGA) (2). Despite these advances, access limitations, cohort imbalance, and domain shift between normal and condition populations remain practical bottlenecks for downstream model development.

Classical upsampling techniques, such as linear interpolation via SMOTE, assume local Euclidean linearity and fail to capture non-linear, tissue-specific co-expression networks. Also, while Generative Adversarial Networks (GANs) offer nonlinear modeling, they are prone to mode collapse in tabular domains.

Generative methods enable the augmentation of scarce clinical datasets while facilitating the analysis of underlying distributional shifts. Diffusion probabilistic models provide stable likelihood-based generative training (3) with strong fidelity under improved schedules and samplers (4, 5). In parallel, JEPA-style representation learning emphasizes predictive latent structure over direct input reconstruction (6), which is attractive for modeling cross-domain biological shifts.

Here we introduce *Hoike*^1^, a hybrid framework that couples a cross-domain JEPA module with latent diffusion to generate condition-specific transcriptomes from normal reference context. The central design goal is to preserve global tissue structure while learning directional transitions toward condition-associated expression regimes.

Recent transcriptomic generative approaches include variational modeling frameworks for expression data (7). *Hoike* differs by explicitly separating reference-context encoding from condition-target encoding, then using diffusion in the learned latent space for sample synthesis.

Compared with purely conditional diffusion over raw gene vectors, *Hoike* conditions on a learned continuous context vector from normal references, providing a biologically anchored bridge between source and target domains. Compared with standalone JEPA predictors, *Hoike* adds iterative denoising to support stochastic generation of diverse synthetic samples.

## Methods

### Data Sources and Preprocessing

*Hoike* is designed for paired-domain training where a normal reference matrix and a condition matrix share a harmonized gene vocabulary. As a reference set of models, normal references are derived from GTEx tissue expression profiles (1), and condition cohorts are prepared from TCGA programs (2).

Input matrices are represented in log_2_(TPM+1) space. The training pipeline aligns overlapping genes between reference and condition tables, filters samples to tissues with available normal baselines, and validates finite non-negative values prior to optimization. These checks are implemented to fail fast on invalid data and to preserve biological interpretability of decoded outputs. An example of the data format is shown in Table 1.

**Table 1.**
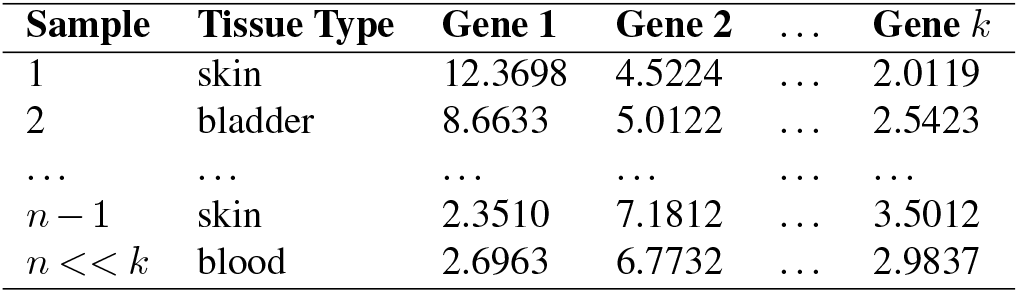
Example data shape used for the input normal and condition samples.

### Problem Formulation

Let *x*_*n*_ ∈ ℝ*G* denote a normalreference expression vector and *x*_*c*_ ∈ ℝ*G* a condition expression vector over *G* shared genes. *Hoike* learns a mapping that preserves tissue-context structure while modeling the condition shift:

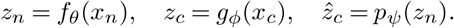

Here, *f*_*θ*_ encodes normal context, *g*_*ϕ*_ encodes condition targets, and *p*_*ψ*_ predicts condition latents from normal context. A diffusion model then learns reverse denoising in latent space conditioned on *z*_*n*_.

### *Hoike* Architecture

The *Hoike* architecture supports a blended JEPA and diffusion-based approach. The model training process accepts a matrix of normal expression values and a condition expression matrix. Both input datasets have the same gene ID columns and a tissue type class column. This architecture is implemented in PyTorch and uses modular engines for the JEPA and diffusion training steps. Data preparation utilities are in `scripts/prepare/`, tensor conversion in `scripts/convert/`, and inference in `scripts/infer/`.

### Cross-Domain JEPA Module

The JEPA component uses asymmetric streams:

1. A baseline encoder *f*_*θ*_ (MLP) maps normal inputs to a continuous context representation.
2. A condition encoder *g*_*ϕ*_ maps condition inputs to target latent vectors.
3. A predictor *p*_*ψ*_ estimates condition latents from normal context.
4. A decoder *d*_*ω*_ maps latent vectors back to geneexpression space.

This design separates contextual anchoring from target-state representation, enabling explicit modeling of normalto-condition transitions.

### Latent Diffusion Module

*Hoike* trains a denoiser *ϵ*_*η*_ over latent vectors using a DDPM-style forward process (3):

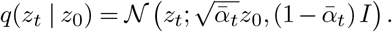

Reverse denoising is conditioned on the JEPA context vector through cross-attention blocks in the latent denoiser. Sampling follows iterative reverse updates, with optional accelerated inference using DDIM (5).

### Optimization Objective

The *Hoike* training objective combines representation alignment and generation constraints. In the implementation used for this repository, optimization uses weighted terms that include:

1. Latent prediction/noise objectives for JEPA and diffusion consistency.
2. Gene-space reconstruction terms through the decoder.
3. Mean/covariance alignment terms to preserve distributional structure.
4. Shift-direction terms encouraging movement from normal toward condition manifolds.
5. Non-negativity penalties to discourage invalid negative expression values after decoding.

Model parameters are optimized with Adam-family optimizers (8).

### Synthetic Generation Procedure

At inference, *Hoike* supports tissue-matched synthetic generation: (1) build a normal context vector for each target condition sample (or from a chosen normal profile); (2) initialize latent noise; (3) perform reverse diffusion conditioned on normal context; (4) decode generated latent vectors to gene-expression profiles; and (5) clamp outputs to non-negative log_2_(TPM+1) values. The full model architecture is shown in Figure 1.

**Fig. 1.**
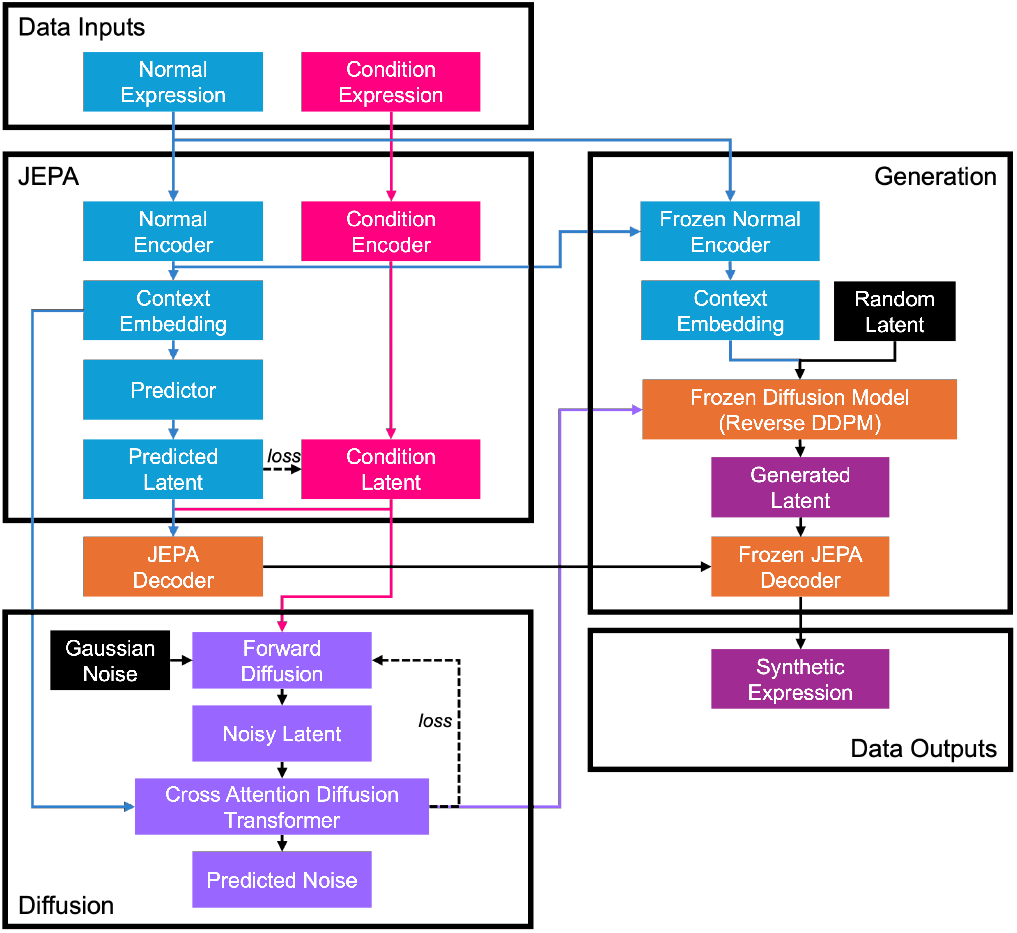
*Hoike* model architecture showing training and generation pathways.

### Loss Weight Customization

A key capability of the *Hoike* framework is that the loss weights (for both the JEPA and diffusion steps) can be customized to fit specific desired characteristics of the synthetic data.

For the JEPA training portion, the loss weights are customizable to guide the semantic learning process of the latent representation of the data. These weights include:

- *latent*: Latent representation loss is primary for JEPA training, as it captures the essential features of the normal-to-condition mapping.
- *gene_recon*: Reconstruction loss on the gene expression space ensures that the predicted condition profiles are biologically plausible and consistent with observed data.
- *mean*: Mean loss encourages the predicted condition profiles to have similar average expression levels to the true condition profiles, promoting overall fidelity.
- *structure*: Structure loss encourages the predicted condition profiles to maintain similar correlation structure to the true condition profiles, preserving biological relationships between genes.
- shift: Shift loss encourages the predicted condition profiles to capture the correct direction of change from normal to condition, ensuring that the model learns the underlying biological shifts.
- *nonneg*: Nonnegativity loss penalizes negative values in the predicted condition profiles, promoting biologically plausible gene expression levels.

Similarly, in the diffusion model training, the loss weights can be customized to guide the learning for denoising the latent representation of the data into realistic gene expression values. There are also weights to promote similar geometries or penalize the generation of negative values, given that the input values are in log_2_(TPM+1) units, which is always *>* 0. These weights are:

- *noise*: Noise prediction loss is the primary objective for training the diffusion model, as it learns to denoise latent representations conditioned on normal states.
- *x0*: X0 reconstruction loss encourages the diffusion model to produce latent representations that can be decoded back into biologically plausible gene expression profiles, ensuring consistency with the original data.
- *gene_recon*: Gene reconstruction loss ensures that the decoded latent representations from the diffusion model produce biologically plausible gene expression profiles, maintaining fidelity to the original data.
- *structure*: Structure loss encourages the decoded latent representations to maintain similar correlation structure to the true condition profiles, preserving biological relationships between genes.
- *nonneg*: Non-negativity loss penalizes negative values in the predicted condition profiles, promoting biologically plausible gene expression levels.

## Evaluation Protocol

The evaluation pipeline compares three distributions: normal vs condition, condition vs synthetic condition, and normal vs synthetic condition. Metrics include:

1. Mean-profile similarity (correlation and MSE).
2. Covariance and geometry distances (Frobenius, affine-invariant, Bures/Wasserstein, Fréchet-type distances) (9).
3. PCA-space centroid and neighborhood comparisons using reference-normal projections.
4. Correlation-structure gaps over high-variance genes.

This triplet protocol quantifies both fidelity to the target condition and retention of biologically coherent structure relative to normal baselines. Full definitions of these metrics are listed in Supplementary Table S1.

## Case Study: Oncological Data

*Hoike* has been tested in this study for multiple tissue-condition pairs in oncology, including GTEx-derived references and TCGA cohorts such as BRCA, GBM, KIRC, LIHC, LUAD, and SKCM. Trained checkpoints are provided for JEPA and diffusion variants under matched tissue settings.

While this manuscript focuses on the framework specification and reproducible implementation details, these cohort-specific quantitative outcomes are to illustrate the performance and utility of *Hoike* on common public datasets.

### Skin Cutaneous Melanoma Example

As an example investigation into the statistical evaluation of the generative realism, we have performed synthetic data generation based on normal skin tissue profiles (from GTEx) and skin cutaneous melanoma condition samples from the TCGA-SKCM project (10).

To evaluate the fidelity and distributional alignment of the synthesized transcriptomic profiles, we established a triplet evaluation protocol comparing three distinct data states: healthy reference baselines (*X*_normal_), true disease targets (*X*_condition_), and model outputs (*X*_synthetic_). Quantitative benchmarks across high-dimensional feature space demonstrate that the *Hoike* framework successfully shifts expression vectors from the normal baseline to the target condition manifold.

Table 2 summarizes the distance metrics and statistical correlations across the benchmark evaluation suite.

**Table 2.** Quantitative similarity and distance metrics across *X, X*, and *X*transcriptomic distributions.

| Metric | Normal vs. Condition | Condition vs. Synthetic | Normal vs. Synthetic | Target Alignment |
| --- | --- | --- | --- | --- |
| Mean Correlation | 0.8669 | **0.9986** | 0.8650 | Achieved |
| Mean MSE | 1.3555 | **0.0159** | 1.3819 | Achieved |
| Covariance MSE | 333,753.50 | **39,790.90** | 81,034.60 | Achieved |
| Frobenius Distance | 14,442.85 | **4,986.92** | 7,116.64 | Achieved |
| Log-Determinant Difference | 22.72 | **243.72** | 253.97 | Achieved |
| Affine-Invariant Distance | 10.74 | **50.46** | 53.89 | Achieved |
| Bures Distance | 13,882.90 | **10,795.25** | 12,219.73 | Achieved |
| Fréchet Distance | 31,064.59 | **10,849.11** | 27,960.25 | Achieved |
| Eigenvalue Correlation | 0.9959 | 0.9661 | **0.9674** | Suboptimal |
| Explained Variance MSE | 0.0006 | **0.0049** | 0.0073 | Achieved |

The primary distance metrics confirm that *X*_synthetic_ aligns closely with *X*_condition_. The Mean Correlation between true condition profiles and synthetic profiles reaches 0.9986, compared to 0.8669 between normal reference and condition samples. Mean Squared Error (MSE) exhibits a significant reduction from 1.3555 (*X*_normal_ vs. *X*_condition_) to 0.0159 (*X*_condition_ vs. *X*_synthetic_). Furthermore, the distance between *X*_normal_ and *X*_synthetic_ (1.3819) closely mirrors the true biological divergence between *X*_normal_ and *X*_condition_ (1.3555), indicating that the model captures the correct trajectory and magnitude of the biological shift rather than over-fitting to the baseline input. This can be seen visually via principal component analysis in Figure 2.

**Fig. 2.**
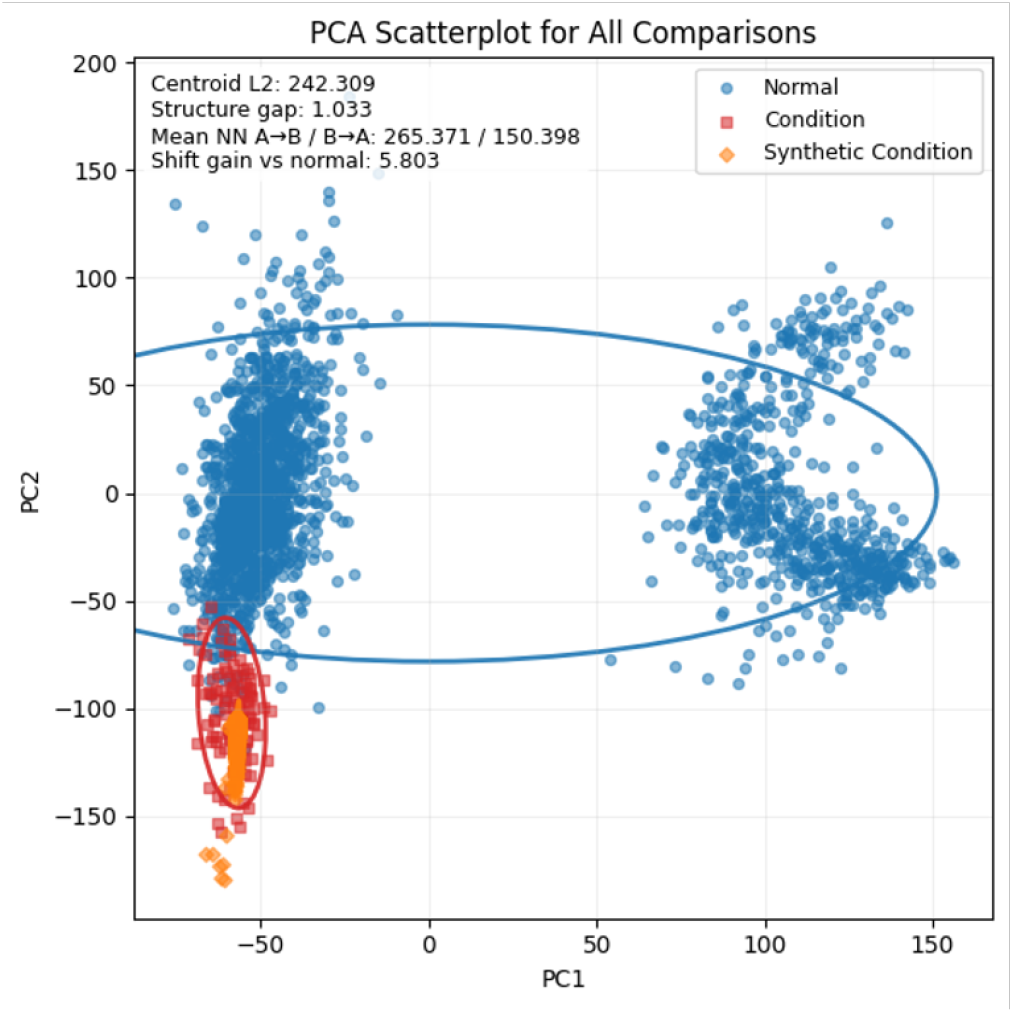
PCA scatterplot showing the positioning of synthetic samples relative to condition SKCM data to normal GTEx data using baseline loss weights.

Results validate the preservation of complex gene coexpression structures. The Covariance MSE decreases by nearly an order of magnitude from 333, 753.50 (*X*_normal_ vs. *X*_condition_) to 39, 790.90 (*X*_condition_ vs. *X*_synthetic_). Similarly, distributional divergence metrics, namely Fréchet Distance (10, 849.11 vs. 31, 064.59) and Frobenius Matrix Distance (4, 986.92 vs. 14, 442.85), demonstrate substantial spatial convergence toward the target disease distribution. While Eigenvalue Correlation shows a minor reduction (0.9661 vs. 0.9959), the discrepancy is minimal and does not detract from the overall structural accuracy of the generated arrays.

### Increasing Variability

If desired, modifying the default loss weights in the training process can increase the variability and shape of the resulting synthetic data. In Figure 2, the normal weights resulted in synthetic samples that were concentrated near the centroid (mean) of the condition samples.

However, if better coverage or more variability is required, loss weights can be modified accordingly.

For example, as shown in Figure 3, the loss weights were modified to increase variability in the synthetic outputs and resulted in a synthetic geometry that better overlaps with the condition data. This modified weighting configuration increased synthetic variance, improving spatial overlap with the target condition manifold at the cost of slight distributional encroachment into the normal tissue space.

**Fig. 3.**
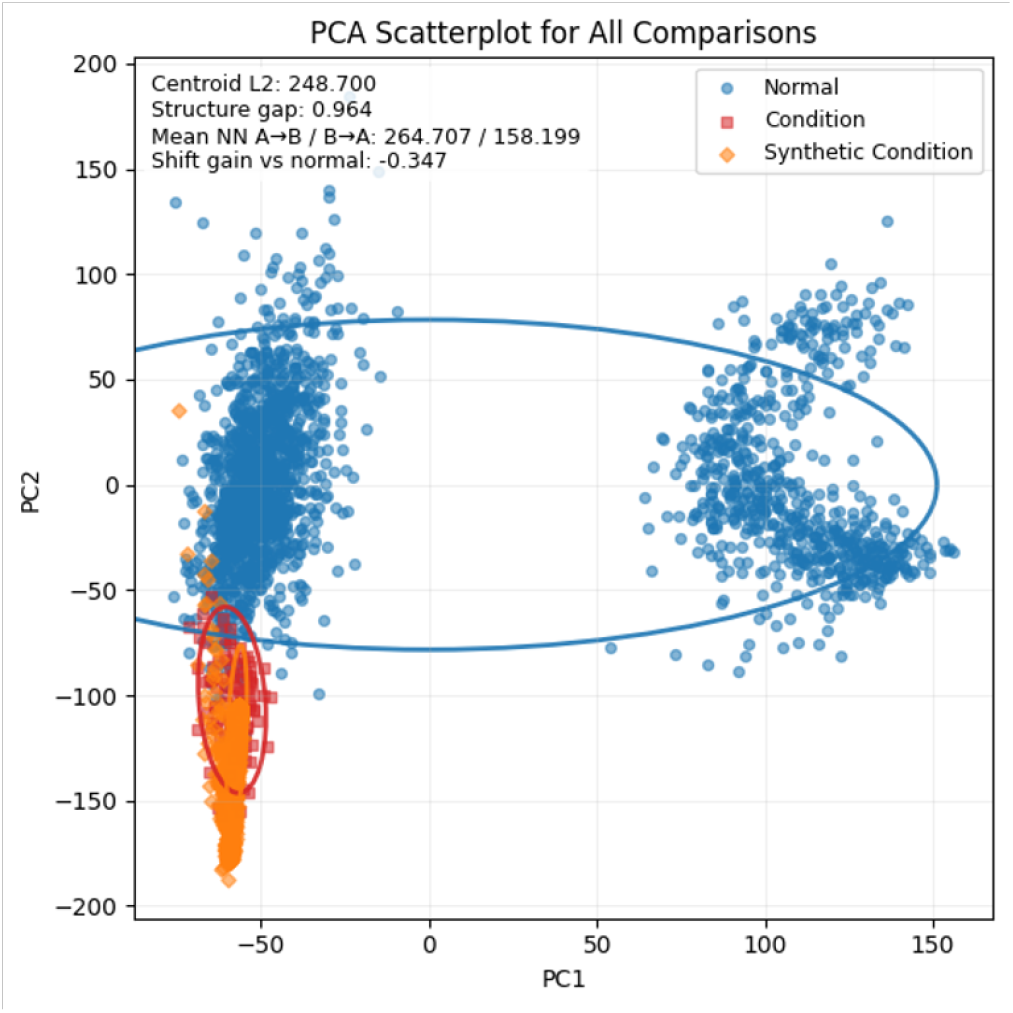
PCA scatterplot showing the positioning of synthetic samples relative to condition SKCM data to normal GTEx data using modified loss weights that increase output variability.

### Cross-Cohort Generative Generalization

To assess generalization across distinct biological contexts, the framework was evaluated across six independent TCGA disease cohorts paired with corresponding GTEx tissue baselines: Glioblastoma Multiforme (GBM), Breast Invasive Carcinoma (BRCA), Kidney Renal Clear Cell Carcinoma (KIRC), Liver Hepatocellular Carcinoma (LIHC), Lung Adenocarcinoma (LUAD), and Skin Cutaneous Melanoma (SKCM). Performance was quantified using two primary geometric metrics:

1. **Centroid Distance (***d*_*C*_): The Euclidean distance between dataset centers of mass in R*G* feature space.
2. **Structure Gap (***S*_*G*_): The structural divergence between normalized gene-to-gene co-expression correlation matrices.

Table 3 presents the cross-cohort generalization benchmarks. Across all evaluated tissue types, the synthetic profiles achieve closer spatial proximity to the true disease centroids than the corresponding healthy baselines (*dC* (*X*condition, *X*synthetic) *< dC* (*X*normal, *X*condition)). The model exhibits high spatial precision in brain (GBM, *d*_*C*_ = 25.80) and lung (LUAD, *d*_*C*_ = 25.53) cohorts despite large baseline shifts (*d*_*C*_ *>* 195). Conversely, cohorts characterized by a higher baseline structural reorganization, such as skin (SKCM, Structure Gap = 0.672), exhibit higher centroid dispersion (*d*_*C*_ = 183.01), indicating increased trajectory variance during continuous cross-attention sampling. Overall, these results confirm that joint-embedding predictive conditioning combined with latent diffusion enables robust crossdomain transcriptomic translation across diverse oncological profiles.

**Table 3.** Multi-tissue generalization evaluation measuring centroid distances (*d*_*C*_) and co-expression structure gaps (*S*_*G*_) across six TCGA cohorts.

| Tissue<br>TCGA ID | Brain<br>GBM | Breast<br>BRCA | Kidney<br>KIRC | Liver<br>LIHC | Lung<br>LUAD | Skin<br>SKCM |
| --- | --- | --- | --- | --- | --- | --- |
| $d_C$ (Cond→Syn) | **25.80** | **29.22** | **139.15** | **68.92** | **25.53** | **183.01** |
| $d_C$ (Norm→Cond) | 261.06 | 192.10 | 172.61 | 182.15 | 195.92 | 151.81 |
| $d_C$ (Norm→Syn) | 245.91 | 182.17 | 143.99 | 140.41 | 193.47 | 290.24 |
| Alignment | ✓ | ✓ | ✓ | ✓ | ✓ | ✓ |
| $S_G$ (Cond→Syn) | 0.599 | 0.785 | 0.729 | 0.904 | 0.787 | 0.539 |
| $S_G$ (Norm→Cond) | 0.413 | 0.486 | 0.375 | 0.298 | 0.270 | 0.672 |
| $S_G$ (Norm→Syn) | 0.701 | 0.783 | 0.742 | 0.908 | 0.863 | 0.855 |

## Discussion

*Hoike* is designed to model condition-associated transcriptomic shifts while retaining normal-context anchoring. The hybrid JEPA-diffusion formulation offers three practical advantages: modular conditioning within a normal biological reference context, the stochastic generation of multiple plausible condition profiles, and the capacity for explicit geometric evaluation of synthetic realism and shift behavior.

By using JEPA, the model processes the relationship between the embeddings (the latent space hidden relationships) rather than the explicit examples. Thus, it appears to learn a more semantic representation of the complex relationships between thousands of genes, allowing the diffusion model to be trained much more quickly and with fewer observations. Limitations currently include dependence on accurate tissue label harmonization, possible sensitivity to cohort preprocessing choices, and the need for careful downstream validation before clinical or mechanistic interpretation. Plus, the weighted loss approach, while it grants control over model outputs, may require trial and error to achieve desired results. Future work includes expanded cross-cohort validation, alternative conditioning schemes (for example, covariates beyond just tissue type), and integration with external perturbation or intervention datasets (including non-oncological data). We wish to also improve the configuration of loss weights, perhaps by providing additional templates for various use cases or automated tuning in a hyperparameter search-like interface.

Although empirical validation has focused primarily on oncological pathologies sourced from The Cancer Genome Atlas (TCGA), the underlying architecture is domain-agnostic. Subsequent evaluations will target non-oncological conditions characterized by subtle, multi-gene expression shifts, including autoimmune diseases, neurodegenerative disorders, and metabolic syndromes. Extending validation to noncancerous and perturbed cohorts will help establish the sensitivity boundaries of the framework without overshooting delicate transcriptional targets.

As can be seen in some of the condition outputs, there are some non-linear or gradient structures in the synthetic samples after PCA comparisons. Future improvements should detect this geometry for retraining with different loss weights. Also, there are improvements to be made in the post-generation data evaluation to better automate the detection of a “good” synthetic data output.

Lastly, synthetic data outputs from models trained with the *Hoike* framework should be validated in differential expression and machine learning analyses to assess the generated data’s effects on the statistics or predictions in translational bioinformatics studies.

While models trained using JEPA-based frameworks have been published for self-supervised learning from images (11), motion dynamics in video (12), and tabular data generation (13), our application of JEPA and diffusion for the generation of gene expression data is quite novel.

In this work, we have shown the utility of *Hoike* in the fast, controllable generation of synthetic data given a combined input of available normal samples and a fewer number of condition samples on which to train a model. This framework helps to address the issue of limited disease-level data in transcriptomics studies and will help to unlock the ability to use that data in impactful clinical research.

## Declaration of Interests

Author CTF is the owner of Tuple, LLC, a biotechnology consulting firm, and its subsidiary, Silico Biosciences. The remaining authors declare that the research was conducted in the absence of any commercial or financial relationships that could be construed as a potential conflict of interest.

## Acknowledgments

We acknowledge the following entities at the University of North Carolina at Charlotte: the Center for Computational Intelligence to Predict Health and Environmental Risks (CI-PHER), the Department of Bioinformatics and Genomics, and the School of Data Science.

We also thank Jonas Elmerraji for his help and ideas surrounding the multidimensional evaluation metrics.

## Code and Data Availability

All code, data, results, and additional analyses are openly available on GitHub at: https://github.com/silicobio/hokie. This repository includes the open-source logic for retrieving and shaping GTEx and TCGA data. Also, there is example training code to create models and generate synthetic datasets.

Model weights for the oncology case study and input GTEx and TCGA datasets are hosted on Hugging Face at https://huggingface.co/collections/silicobio/hoike.

## Funding Statement

Funding for cloud computational resources was provided by the Microsoft Most Valuable Professionals program.

## Supplementary Materials

**Supplementary Table S1.**
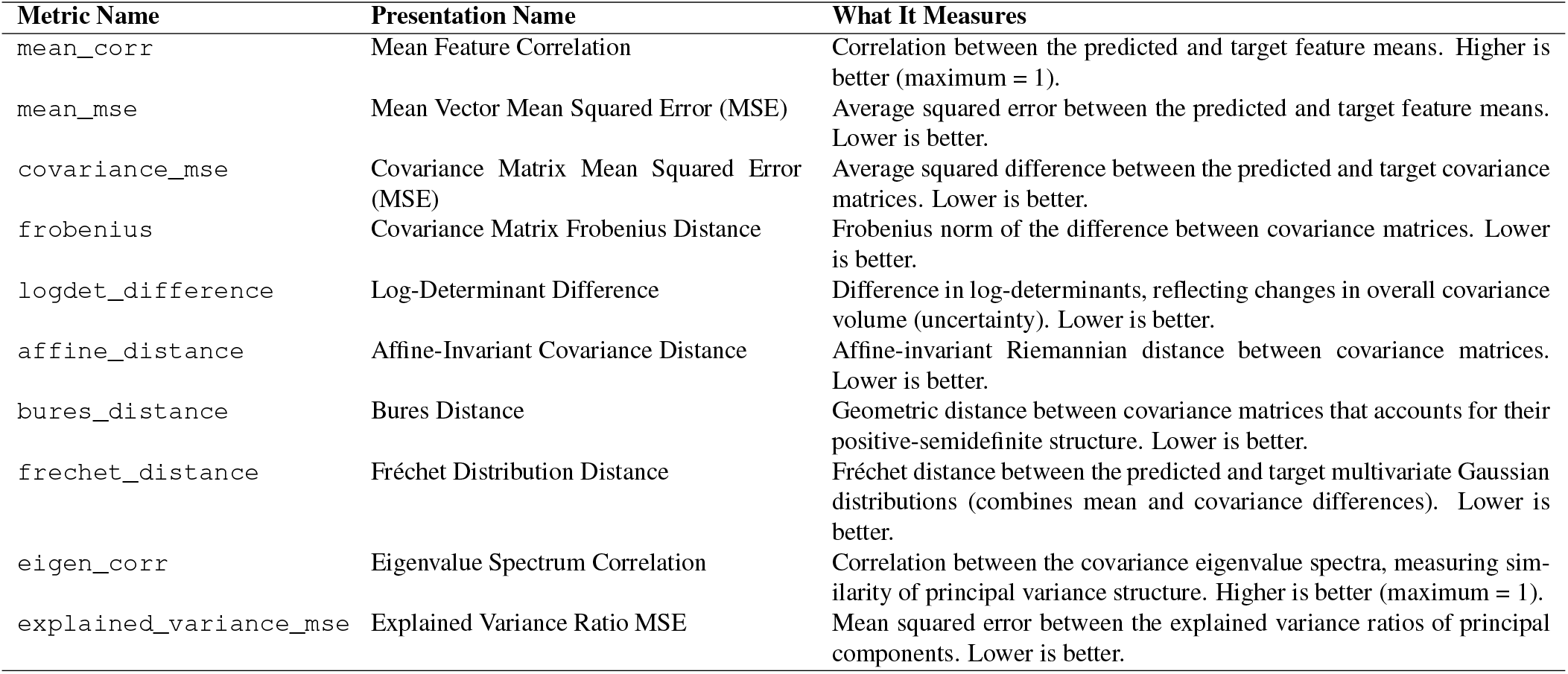
Summary of evaluation metrics used for comparing predicted and target feature distributions.

## Footnotes

1 *Hoike* means “to express” in native Hawaiian.

## Notes

https://github.com/silicobio/hoike

https://huggingface.co/collections/silicobio/hoike

